# Global abundance patterns, diversity, and ecology of Patescibacteria in wastewater treatment plants

**DOI:** 10.1101/2023.10.25.562895

**Authors:** Huifeng Hu, Jannie Munk Kristensen, Craig William Herbold, Petra Pjevac, Katharina Kitzinger, Bela Hausmann, Morten Kam Dahl Dueholm, Per Halkjaer Nielsen, Michael Wagner

**Affiliations:** Centre for Microbiology and Environmental Systems Science, University of Vienna. Djerassiplatz 1, 1030 Vienna, Austria; Center for Microbial Communities, Department of Chemistry and Bioscience, Aalborg University, Aalborg, Denmark; Joint Microbiome Facility of the Medical University of Vienna and the University of Vienna, Vienna, Austria; Division of Clinical Microbiology, Department of Laboratory Medicine, Medical University of Vienna, Vienna, Austria; Doctoral School in Microbiology and Environmental Science, University of Vienna. Universitätsring 1, 1010 Vienna, Austria; Te Kura Pūtaiao Koiora | School of Biological Sciences, Te Whare Wānanga o Waitaha | University of Canterbury, Christchurch, New Zealand

**Keywords:** Patescibacteria, wastewater treatment plants, 16S rRNA gene amplicon sequencing, diversity, network analysis, host prediction

## Abstract

**Background:** Microorganisms are responsible for nutrient removal and resource recovery in wastewater treatment plants (WWTPs), and their diversity is often studied by 16S rRNA gene amplicon sequencing. However, this approach underestimates the abundance and diversity of Patescibacteria due to the low coverage of commonly used PCR primers for this highly divergent bacterial phylum. Therefore, our current understanding of the global diversity, distribution, and ecological role of Patescibacteria in WWTPs is very incomplete. This is particularly relevant as Patescibacteria are considered to be associated with microbial host cells and can therefore influence the abundance and temporal variability of other microbial groups that are important for WWTP functioning.

**Results:** Here, we evaluated the *in silico* coverage of widely used 16S rRNA gene-targeted primer pairs and redesigned a primer pair targeting the V4 region of bacterial and archaeal 16S rRNA genes to expand its coverage for Patescibacteria. We then experimentally evaluated and compared the performance of the original and modified V4-targeted primers on 565 WWTP samples from the MiDAS global sample collection. Using the modified primer pair, the percentage of ASVs classified as Patescibacteria increased from 5.9% to 23.8%, and the number of detected patescibacterial genera increased from 560 to 1,576, while the detected diversity of the remaining microbial community remained similar. Due to this significantly improved coverage of Patescibacteria, we identified 23 core genera of Patescibacteria in WWTPs and described the global distribution pattern of these unusual microbes in these systems. Finally, correlation network analysis revealed potential host organisms that might be associated with Patescibacteria in WWTPs. Interestingly, strong indications were found for an association between Patescibacteria of the Saccharimonadia and globally abundant polyphosphate-accumulating organisms of the genus *Ca.* Phosporibacter.

**Conclusions:** Our study (i) provides an improved 16S rRNA gene V4 region-targeted amplicon primer pair inclusive of Patescibacteria with little impact on the detection of other taxa, (ii) reveals the diversity and distribution patterns of Patescibacteria in WWTPs on a global scale, and (iii) provides new insights into the ecological role and potential hosts of Patescibacteria in WWTPs.

## Background

Wastewater treatment is currently the largest industrial application of biotechnology in terms of volume, and is becoming increasingly important due to population growth and the urgency of environmental protection. Globally, activated sludge systems are the most widely applied wastewater treatment systems. In activated sludge, and other types of WWTPs, microbial communities play crucial roles in wastewater processing, nutrient removal, and bioenergy production [1]. 16S ribosomal RNA (rRNA) gene amplicon sequencing is commonly used to study the microbial communities of these engineered ecosystems [2–4]. However, there are two major challenges with this method: (i) the lack of complete reference databases that can provide a comprehensive taxonomic classification for many uncharacterized environmental microbes, and (ii) that widely used general primer sets do not adequately cover some important lineages within the tree of life. To overcome the first challenge, a comprehensive ecosystem-specific database including more than 80,000 full-length 16S rRNA gene sequences was recently built from a large set of WWTPs samples collected worldwide (Microbial database for activated sludge, MiDAS), facilitating domain-to-species level taxonomic classification for amplicon-based WWTP studies [5]. To solve the second challenge, several studies have reported modifications of primers to improve coverage against specific lineages [6–8], but there are still some lineages with insufficient primer coverage, which can lead researchers to overlook their distribution and importance in various environments, including activated sludge.

One widespread microbial group often incompletely covered by amplicon sequencing is Patescibacteria, also referred to as Candidate Phyla Radiation (CPR) [9]. Patescibacteria are a recently discovered lineage that is widespread in various environments, including groundwater [10,11], freshwater [12,13], the human oral cavity [14], and also WWTPs [5,15,16]. The term CPR was initially proposed for these organisms [10], and they were defined as a superphylum radiation that contains more than 74 phyla for which the branching order couldn’t be accurately determined [17]. Recently, the Genome Taxonomy Database (GTDB) taxonomy [18] amalgamated CPR into a single phylum named Patescibacteria based on genomic evidence. In this manuscript, we have opted for using the genome-based taxonomy, and will refer to this group of microorganisms as Patescibacteria.

Patescibacteria are characterised by divergent ribosomal RNA (rRNA) genes and peculiar ribosome structure, limited metabolic capacity, a reduced genome size, and small cell size [10,19] The compositional and structural peculiarities of patescibacterial ribosomes include the absence of ribosomal protein L30 (rpL30) in all known genomes, and the absence of rpL9 and rpL1 in several groups, including the classes Microgenomatia and Saccharimonadia (also known as Saccharibacteria/TM7) [10]. The presence of self-splicing introns in 16S and 23S rRNA genes has also been reported for Patescibacteria. Such introns in the 16S rRNA gene can be located at multiple positions within the gene and can be >5 kb in length [10].

Phylogenetic analyses of the 16S rRNA gene and concatenated ribosomal proteins both show that Patescibacteria encompassed a huge diversity and suggest a rapid evolutionary rate in this group of bacteria [17]. Genome-based analyses have predicted that Patescibacteria lead a symbiotic lifestyle. Consistently, several members of Patescibacteria have been shown to associate with different hosts, including eukaryotes, archaea, and bacteria. A member of the Parcubacteria (Paceibacteria in the GTDB) was reported to be a protist endosymbiont (*Paramecium bursaria*) [13]. Another symbiotic relationship was found between an archaeon (*Methanothrix*) and another member of the Paceibacteria [20]. Saccharimonadia are the patescibacterial group that has been most extensively studied in terms of host association. To date, representatives of three genera of the Saccharimonadia have been co-isolated together with bacterial hosts from the human oral cavity, WWTPs, and insects, respectively, and all have been shown to have an epiparasitic lifestyle with hosts from the phylum Actinomycetota [14,21,22]. However, the lifestyle and ecology of the majority of Patescibacteria remain unknown.

While metagenomic studies are becoming more common, the research of low abundant organisms, such as members of the Patescibacteria, still relies on amplicon sequencing-based approaches. Consequently, several studies have developed or modified 16S rRNA gene-targeted primer pairs to improve coverage of Patescibacteria or other groups poorly covered by existing primers. However, these studies have either focused on specific lineages or on specific habitats [6–8]. Thus, to date, no published primer pair covers the whole patescibacterial group. Here, we modified a commonly used universal 16S rRNA gene-targeted primer pair to significantly improve its coverage for all known patescibacterial groups, and applied the newly developed primers to study the diversity, global distribution and potential hosts of Patescibacteria in 565 WWTP samples from around the world. Although we examined the primers coverage against Patescibacteria in WWTP samples, the theoretical coverage is also much enhanced for non-WWTP Patescibacteria.

## Results

### Insufficient coverage of Patescibacteria by commonly used 16S rRNA gene primer pairs and design of a new primer pair

*In silico* coverage data of the three most widely used primer pairs for 16S rRNA gene amplicon sequencing **(Table S1)**, targeting the V1-V3, V3-V4, or V4 hypervariable region, were compared for nineteen bacterial phyla and one archaeal phylum with the highest database representation. In these analyses, we focused on perfect matches (0 mismatches) between primers and 16S rRNA gene sequences in the SILVA SSU rRNA gene reference database [23] and the WWTP-specific MiDAS 16S rRNA gene database [5], but also included a coverage analysis allowing for a single mismatch with one of the two primers **(Table S2-S5).**

We first evaluated the coverage of these primers against the SILVA SSU rRNA gene database. As expected, the V1-V3 and V3-V4 primers showed low coverage of archaea, since these primers were designed to target only bacterial 16S rRNA genes. The V1-V3 primer pair had the lowest overall *in silico* coverage, with less than 50% cumulative coverage across all bacterial phyla and only 5.3% for all Patescibacteria. The V4 primer pair, designed to target both bacterial and archaeal 16S rRNA genes, had the best overall coverage. Over 85% of sequences from most phyla showed no mismatches to this primer pair, but it still covered only 19.6% of Patescibacteria. The V3-V4 primer pair showed better but still only moderate coverage of Patescibacteria (57.6%) and much poorer coverage of some other bacterial phyla (e.g. Chloroflexi (40.8%) and Armatimonadota (31.5%)) than the V4 primers **(Table S2)**. Within the Patescibacteria, both the V3-V4 and V4 primer pairs had mismatches to nearly all Microgenomatia and Dojkabacteria sequences. The V4 primer pair also showed low coverage of Saccharimonadia (4.7%), Parcubacteria (6.3%), ABY1 (0.5%) and WWE3 (0%). The V3-V4 primer pair showed better coverage of ABY1 (75.6%), Parcubacteria (55.6%) and Saccharimonadia (84.8%) than the V1-V3 primer and V4 primer pairs **(Table S3).**

*In silico* coverage of the 16S rRNA gene sequences from the MiDAS 4.8.1 database was also evaluated for the V3-V4 and V4 primer pairs. The V1-V3 primer pair (27F/534R) could not be evaluated against MiDAS 4.8.1 because sequences from this database were trimmed after the 27F primer binding site (Dueholm et al., 2020). For sequences included in MiDAS 4.8.1, the V4 primer set showed >70% coverage of the 20 most represented phyla, except for Patescibacteria (15.2%) and Chloroflexi (60.9%) **(Table S4)**. For Patescibacteria, the V4 primer pair targeted only 8% of the Saccharimonadia, 0.5% of the Microgenomatia, 0.2% of the ABY1, and 0% of the Parcubacteria 16S rRNA gene sequences in MiDAS 4.8.1, whereas it covered 93.4% of gracilibacterial 16S rRNA gene sequences retrieved from WWTP systems. The V3-V4 primers showed better coverage of the WWTP Patescibacteria (67.0%) than the V4 primers, covering more Saccharimonadia (88.8%), Parcubacteria (71.8%), ABY1 (87.2%), and a similar fraction of Gracilibacteria (84.0%) 16S rRNA gene sequences **(Table S5).** But consistent with the results obtained using the SILVA database, the V3-V4 primer pair showed poorer coverage of other, non-patescibacterial phyla in the WWTP database.

We additionally evaluated a recently published V4-V5 primer pair (515Yp-min/926Rp-min), modified specifically to capture patescibacterial 16S rRNA gene sequence diversity in marine environments [8,24] using the SILVA and MiDAS 4.8.1 databases. While being well suited for the analysis of Patescibacteria in marine samples, this primer pair shows only overall moderate coverage (37.1%) of Patescibacteria in the SILVA database, and likewise only a moderate coverage of 30.8% of Patescibacteria sequences in the WWTP-specific MiDAS 4.8.1 database **(Table S4)**.

Based on the high general coverage of bacterial and archaeal 16S rRNA gene sequences, and the high overall coverage of WWTP derived 16S rRNA gene sequences by the V4 primer pair (515F-806R) in the *in silico* analysis, this primer pair was chosen for modification to increase its coverage of Patescibacteria. Another reason for choosing the V4 primer pair for further improvement was that this primer pair is predicted to produce shorter amplicons than the V3-V4 primers. Currently, most 16S rRNA gene amplicon studies use short-read Illumina platforms to generate sequence data, which balances quality and expense to produce useful community profiles [25]. Shorter amplicons have also been shown to recover community structure more accurately than longer amplicons [26].

The original V4 primer pair was modified by adding degeneracy bases at 5 positions (8/11/12/13/18) of the forward primer (515F) and 4 positions (1/7/10/14) of the reverse primer (806R). Additionally, we changed the 9th position of the forward primer from M to A **(Table S6)**. These modifications of the V4 primer set resulted in a significantly improved full-match *in silico* coverage against the SILVA database for Chloroflexi (from 56.7% to 76.0%), Spirochaetota (from 71.7% to 79.6%), and Patescibacteria (from 19.6% to 88.9%). Notably, coverage was strongly increased for several Patescibacteria classes including the Microgenomatia (90.9%), Parcubacteria (80.8%) and Dojkabacteria (85.5%) which were poorly covered (1.2%, 6.3% and 0%, respectively) with the original primers. However, due to the change in the ninth position in the original forward primer, the *in silico* coverage of the modified primer set for the Euryarchaeota decreased **(Table S2; Table S4).**

### A global collection of waste water treatment plant samples

The samples **(Figure 1)** and metadata **(Table S7)** used for our analysis were collected by the MiDAS global consortium. This sample set represented 565 WWTPs from 6 continents, 30 countries and 380 cities. Most of the samples (448/79.3%) were from activated sludge plants, but samples from biofilters, moving bed bioreactors (MBBR), membrane bioreactors (MBR), and granular sludge were also included **(Figure 1)**. Among the activated sludge samples, most (n=194) were from plants designed for carbon removal with nitrification and denitrification (C, N, DN; 43.3%), 92 were from plants designed for carbon removal only (C; 20.5%), 90 were from plants designed for carbon removal with nitrogen and enhanced biological phosphorus removal (C, N, DN, P; 20.1%), and 46 were from plants designed for carbon removal with nitrification (C, N; 10.3%).

**Figure 1.**
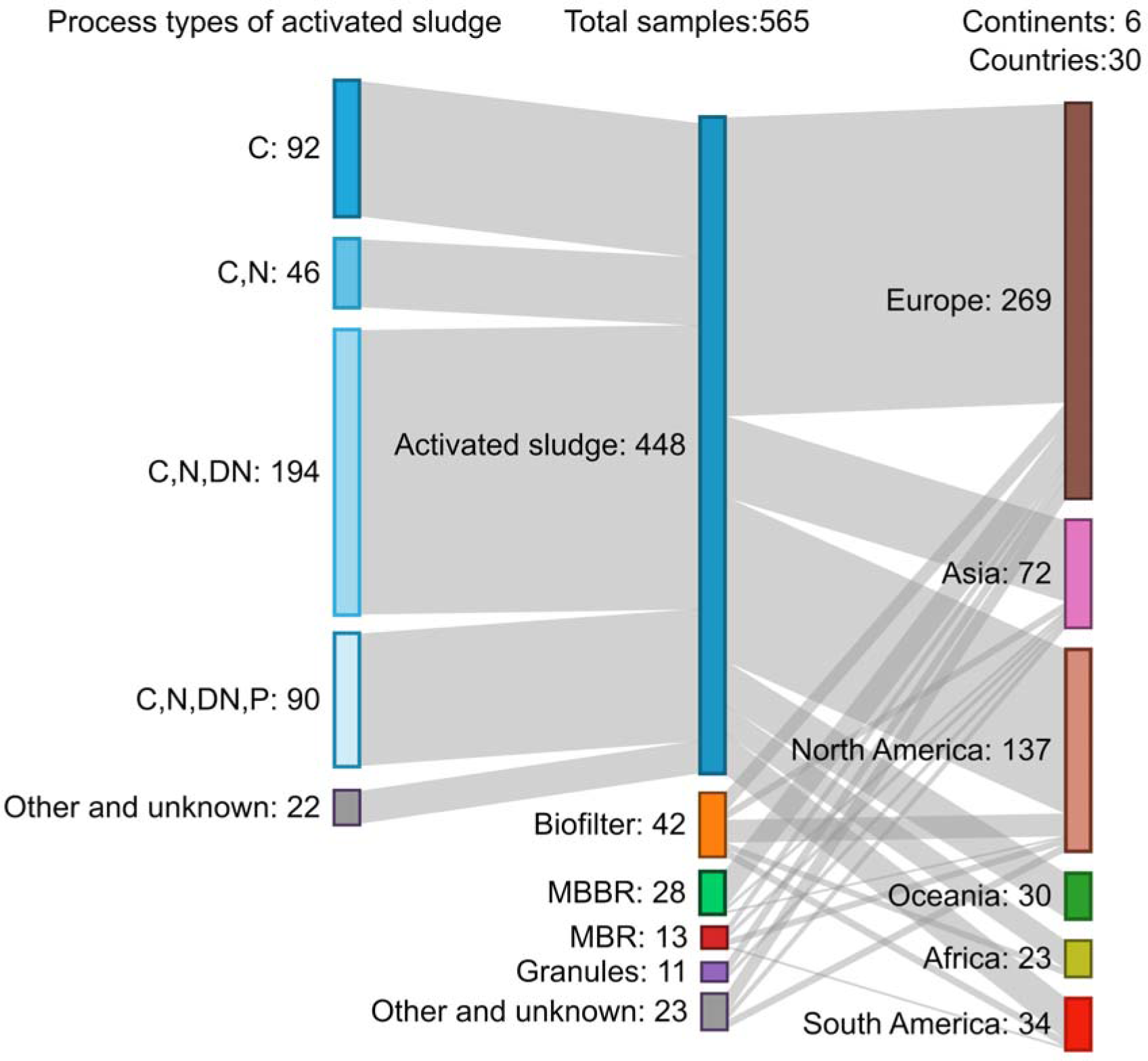
Global WWTP sample set used in this study. Sankey diagram showing the geographical source, plant types and activated sludge process types of 565 analysed WWTP samples. MBBR: moving bed bioreactor; MBR: membrane bioreactor; C: carbon removal; C,N: carbon removal with nitrification; C,N,DN: carbon removal with nitrification and denitrification; C,N,DN,P: carbon removal with nitrogen removal and enhanced biological phosphorus removal (EBPR).

### Modified primers reveal a more complete picture of Patescibacteria diversity and abundance in wastewater treatment plants

Consistent with the *in silico* primer coverage predictions, 16S rRNA gene amplicon datasets generated from the global activated sludge sample collection using our modified V4 primers detected significantly higher Patescibacteria ASV richness and diversity than 16S rRNA gene sequencing datasets generated with the original V4 primers **(Figure 2 a-d)**. At the phylum level, the average relative abundance of Patescibacteria in the activated sludge samples increased dramatically from 1.5% ± 1.4% (mean ± sd) to 18.5% ± 11.1% **(Supplementary Figure 1)**, and the cumulative number of patescibacterial ASVs increased from 5.9% to 23.8% across all samples. The diversity of the total microbial communities in datasets generated using the modified V4 primers was not significantly different compared to datasets obtained using the original V4 primers **(Figure 2 e-f)**. After removing reads classified as Patescibacteria, a linear regression was performed comparing the ASV-based richness of non-patescibacterial taxa detected in each sample by the modified and original primer sets, showing a slightly decreased richness detected by the modified primer sets **(Supplementary Figure 2)**. We performed a phylum level regression analysis to investigate how the ASV richness of non-patescibacterial phyla was affected by the primer modification. Most phyla showed only a slight difference in observed richness, with a predicted slope of ASV richness of original vs. modified primers between 0.6 and 1.6 **(Supplementary Figure 3)**. However, as expected, Euryarchaeota were strongly underestimated by the modified primers, which only detected 15% of the ASV richness detected by the original primers, likely due to the change from M(A/C) to A in the modified forward primer **(Table S6)**. On the other hand, we observed an increased richness of several non-patescibacterial phyla, i.e. Chloroflexi and eight additional phyla in the dataset obtained with the modified primers **(Supplementary Figure 3).**

**Figure 2.**
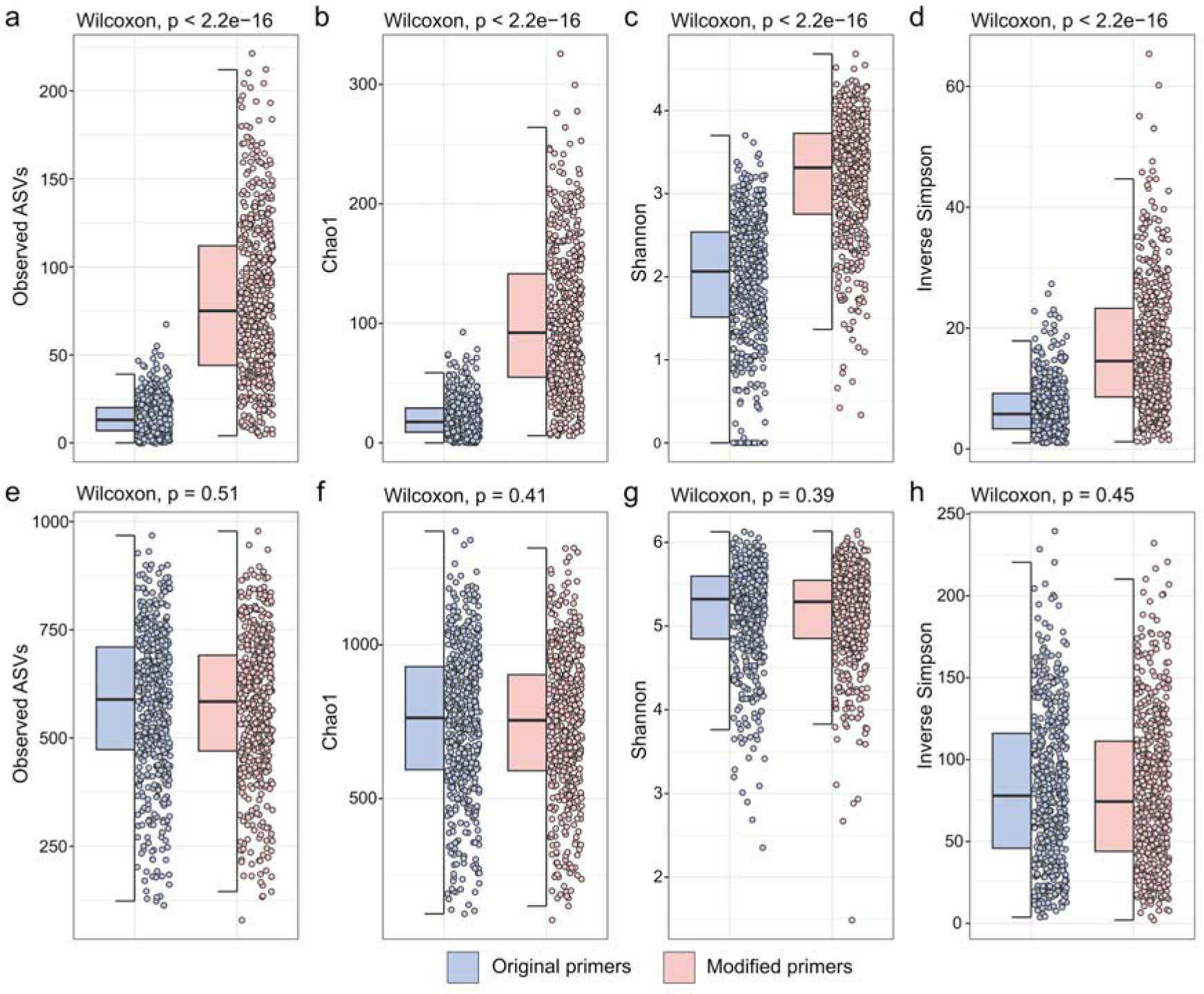
A comparison of patescibacterial and total microbial diversity captured by both primer pairs. Alpha diversity indices (ASV richness, chao1 index, Shannon index and inverse Simpson index) of the Patescibacteria (a-d) and the total microbial community (including Patescibacteria) (e-h) detected by the original primer and the modified primer pairs. Wilcoxon rank sum tests were used to test for significant differences between two groups.

Next, the ASV-based richness within each genus with >0.01% average relative abundance across all samples detected by both the original and modified primer pairs was examined to determine whether the modified V4 primers systematically detected more or less intra-genus diversity than the original primers. It is important to note that the term “genus” used in our study refers to a 94.5% 16S rRNA gene sequence identity threshold [27,28]. Higher ASV richness of some non-patescibacterial genera was detected by the modified primers, for example a 4.27 times higher ASV richness for the genus *Neochlamydia* within the phylum Verrucomicrobiota or a 2.45 higher ASV richness for the genus *Ca.* Villigracilis within the phylum Chloroflexi. The phylum Chloroflexi encompasses most genera (n=29) for which we detected a higher ASV richness with the modified primers **(Table S8).**

### Novel genus-level diversity of Patescibacteria in wastewater treatment plants detected by the modified V4 targeted primers

To compare the genus level diversity of Patescibacteria detected with the two V4 targeted primer pair versions, we selected genera that were detected with at least one primer pair version in at least one sample with more than 0.1% relative abundance. Genera detected at a lower relative abundance were excluded from this analysis. In total, we detected ASV affiliated with 959 patescibacterial genera with the modified primers and 224 genera with both primers. ASVs affiliated with 7 genera of Patescibacteria were only detected in amplicon datasets generated with the original primers. These 7 genera reached a maximum cumulative relative abundance of 0.005% of the total community and 0.39% of Patescibacteria in the dataset generated with the original primer pair. Differential abundance analysis of ASVs affiliated with the 224 genera detected with both primer pairs resulted in 62 (27.7%) genera being significantly more relatively abundant in datasets generated with the modified primer pair (Wald test p-adj< 0.05) **(Figure 3).**

**Figure 3.**
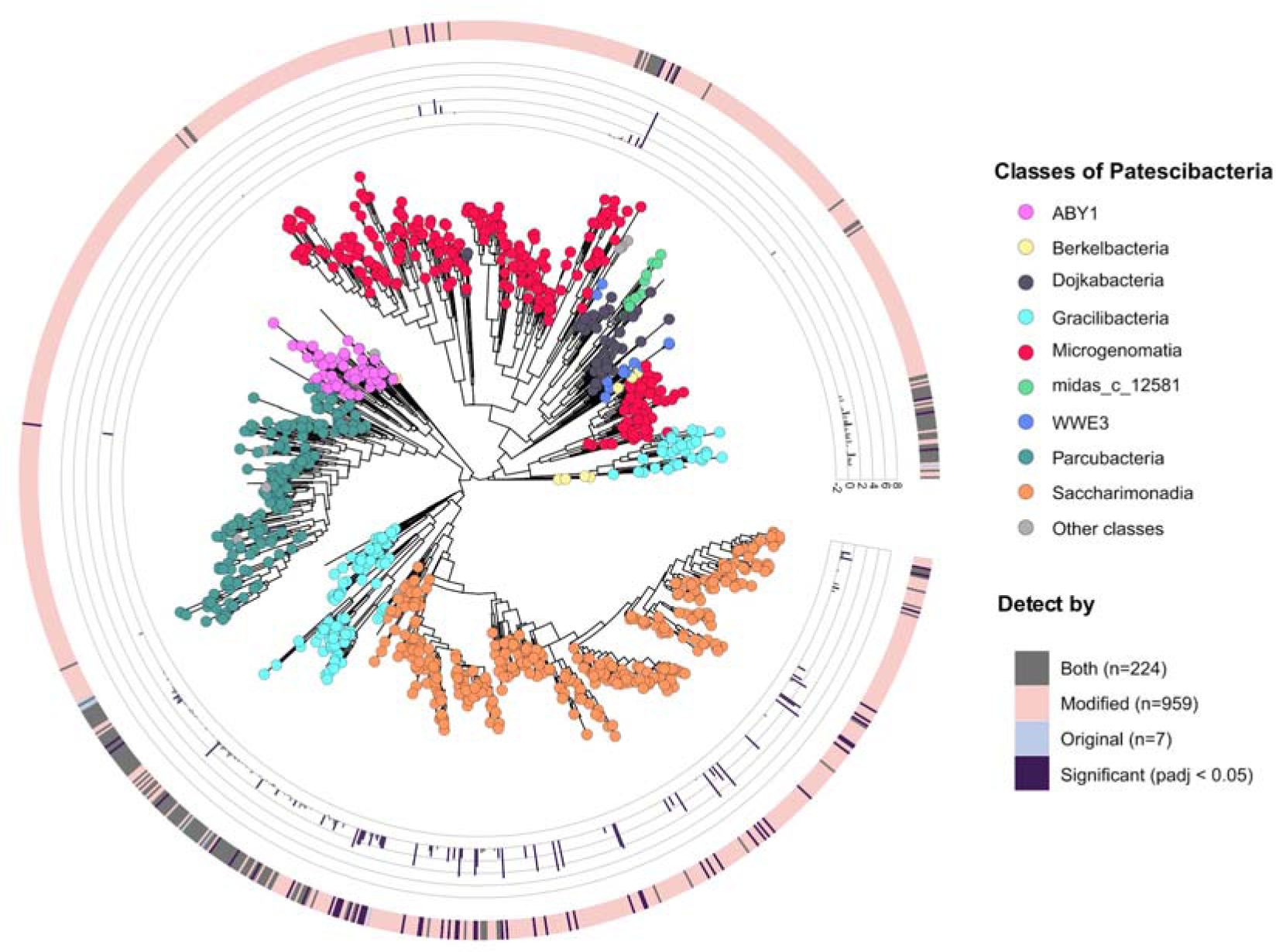
Phylogenetic tree of representative ASVs of patescibacterial genera detected in the global WWTP sample set. The tree includes representative ASVs of genera which were detected in at least one sample with more than 0.1% relative abundance by at least one V4 targeted primer pair version. The nine main patescibacterial classes are depicted by differently colored circles on the tree. For all ASVs detected by both primer sets, the bar plot shows the log fold change (LFC) of read abundance of original vs. modified primer sets. The color of the outermost band represents the detection of each genus by only the original (light blue), only the modified (light pink), or both primers (grey). Dark purple (in both color band and barplot) indicates genera, where the modified primer set detected significantly higher abundances compared to the original primers (adjusted p-value<0.05 calculated by the Wald test).

### Novel amplicon sequence variants revealed by the modified primer pair

16S rRNA genes of Patescibacteria are highly divergent and have thus far likely been undersampled by amplicon studies [10]. To evaluate whether the modified V4 primers enable the detection of previously unknown lineages of Patescibacteria, we determined the proportion of ASVs from each sample that could not be mapped to existing databases (MiDAS 4.8.1 and SILVA r138.1).

We found that within individual samples, significantly more patescibacterial ASVs generated with the modified primers (25.0% ± 11.5%) could not be mapped to the MiDAS database with a high identity (>99%), when compared to patescibacterial ASVs generated with the original primers (17.5% ± 13.9%), which reflects a higher level of phylogenetic novelty detected by the modified primers **(Figure 4a).** It is also noteworthy that as many as 83.1% ± 7.4% of patescibacterial ASVs within individual samples generated by the modified primers and 59.6% ± 16.8% generated by the original primers could not be mapped as high identity hits to the SILVA database **(Figure 4c)**. For non-Patescibacteria ASVs, a similar fraction of ASV obtained with the modified primer pair and the original primer pair within individual samples could be mapped to the two databases **(Figure 4b, 4d)**. Cumulatively, out of all the 9,197 patescibacterial ASVs detected with the modified primers, 4,927 (53.5%) could not be mapped to the MiDAS database with a high identity (>99%), while this was the case for only 655 (36.1%) of the 1,816 patescibacterial ASVs generated with the original primers. For the SILVA database, as many as 8,221 (89.3%) and 1,501 (82.7%) of the patescibacterial ASVs generated with the modified and original primers, respectively, could not be mapped with a high identity (>99%). These higher cumulative unmapped patescibacterial ASV fractions compared to individual sample-based unmapped ASV fractions result from the higher prevalence of ASVs with high identity hits in both databases across all samples (**Supplementary Figure 4).**

**Figure 4.**
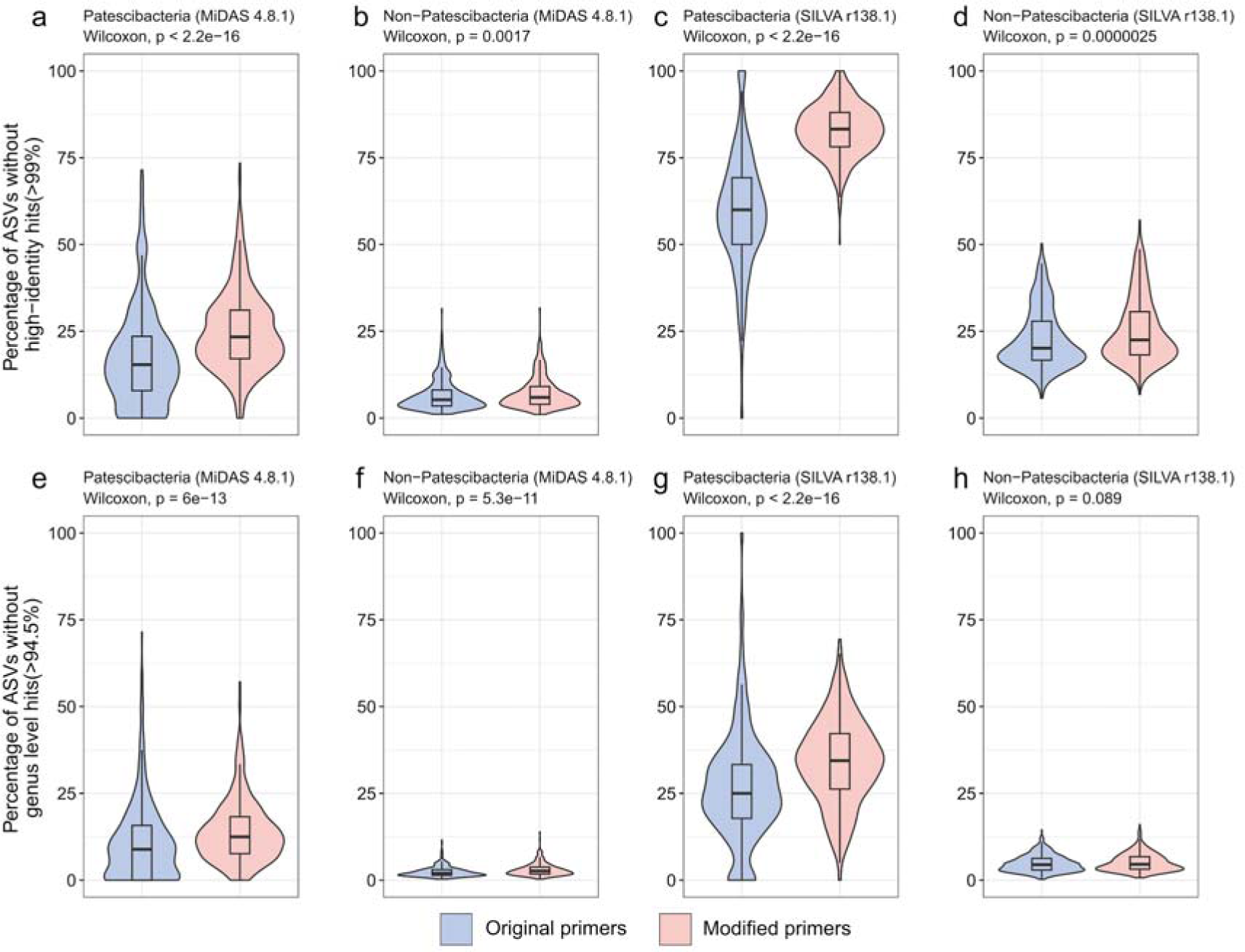
Sequence novelty comparison between 16S rRNA gene amplicon sequence datasets generated by both primer pairs. Percentage of patescibacterial and non-patescibacterial ASVs without high identity hits (99%) in each sample against the MiDAS 4.8.1 database and the SILVA r138.1 database (a-d). Percentage of patescibacterial and non-patescibacterial ASVs without genus level identity hits (94.5%) in each sample at genus level against the MiDAS 4.8.1 database and the SILVA r138.1 database (e-f).

Then, we compared the percentage of ASVs within individual samples generated with the modified primer pair that could not be mapped to the MiDAS 4.8.1 and the SILVA r138.1 database at the genus level threshold (94.5%). Using the MiDAS 4.8.1 database and taxonomy framework, 14.0% ± 9.0% of Patescibacteria ASVs and 3.0% ± 1.8% of non-Patescibacteria ASVs could not be mapped at the genus level **(Figure 4e, 4f)**. To the SILVA database, 34.1% ± 12.2% of Patescibacteria ASVs and 5.1% ± 2.6% of non-Patescibacteria ASVs in each individual sample could not be mapped at the genus level **(Figure 4g, 4h)**. We further explored the taxonomic affiliation of ASVs that could not be classified at genus level by the MiDAS 4.8.1 database. Most of these ASVs were from the classes Microgenomatia (n=837), Parcubacteria (n=472) and Saccharimonadia (n=188). Furthermore, 260 patescibacterial ASVs that could not be classified at class level (in the MiDAS 4.8.1 database) were also detected **(Supplementary Figure 5).** At the genus level threshold (94.5%), 2,868 (31.1%) of all patescibacterial ASVs could not be mapped to the MiDAS database, while this was the case for only 3,345 (11.3%) of all 29,430 non-patescibacterial ASVs. For the SILVA database, 4,966 (53.9%) of the patescibacterial and 4,339 (14.7%) of the non-patescibacterial ASVs could not be mapped at the genus level (**Supplementary Figure 4)**.

Introns in the 16S rRNA gene are frequently detected in Patescibacteria, but do not frequently occur in the V4 hypervariable region [10]. Yet, using the modified V4 primers we detected 17 ASVs with ≥300 bp amplicons that were classified as Patescibacteria, and found that three of those ASVs can be fully mapped to MAGs (metagenome-assembled genomes) from Danish WWTPs [29] **(Table S9).**

### Core genera of Patescibacteria in wastewater treatment plants

Core taxa are characteristic microbial community members of a given environment. Identifying and characterising such taxa is essential for a comprehensive understanding of the microbiology of a system. For WWTPs, core community members have been defined using a relative abundance threshold of 0.1% and a set of prevalence thresholds [30]. According to this classification system, “strict core” community members occur in more than 80% of WWTPs, “general core” members in 50%, and “loose core” members in 20% of the plants [30]. Activated sludge systems harbor a notable loose core community of genera that occur in at least 20% of all plants, constituting more than 50% of the reads in 16S rRNA gene amplicon datasets [5]. In addition to core genera which are globally prevalent, conditionally rare and abundant taxa (CRAT) describe genera that exist in at least one sample with >1% relative abundance, but are not part of the core taxa. Combined, core and CRAT genera can constitute up to 80% of the reads in amplicon datasets from WWTPs [5]. Given the results of the *in silico* primer coverage analysis presented above, the prevalence and relative abundance of Patescibacteria in WWTP has likely been significantly underestimated in previous studies. Consistently, no patescibacterial genera were previously described to be part of WWTP core communities [5].

We compared the richness and relative abundance of Patescibacteria in the 16S rRNA gene amplicon dataset generated with the modified V4 primers across different plant types and we found the activated sludge systems shared similar Patescibacteria abundance and richness with other plant types except biofilters **(Supplementary Figure 6)**. For identification of patescibacterial core genera, we focused on the activated sludge system because it was the most well-sampled system. We identified 23 patescibacterial genera in the four main process types of activated sludge systems from the global WWTP sample set, which were detected in at least 20% of samples with >0.1% relative abundance, fitting the definition of “loose” core taxa **(Figure 5)**. In addition to their frequent occurrence across activated sludge plants, these genera show a global distribution **(Supplementary Figure 7)**. Among these genera, midas_g_2215 is the most abundant genus by average relative abundance (0.72% average, 15.27% highest abundance), followed by the genera midas_g_67 (0.48% average, 9.26% highest abundance), midas_g_4375 (0.37% average, 11.09% highest abundance), midas_g_363 (0.36% average, 7.12% highest abundance) and *Ca.* Saccharimonas (0.35% average, 8.46% highest abundance). Interestingly, only two of the 23 loose core genera have a given genus name (*Ca*. Saccharimonas and TM7a/*Ca*. Mycosynbacter). Members of the genus *Ca*. Saccharimonas are known for their fermentative sugar metabolism [32]. Recently, a member of TM7a has been characterised as a predator of *Gordonia*, which is an infamous foam producer in WWTPs worldwide [21]. Using the GTDB database (r214) as well as metagenomic datasets from Danish WWTPs ([29], we assigned 16S rRNA genes from all MAGs in these two datasets to a MiDAS taxonomy to evaluate the MAGs representation of the core WWTP taxa of Patescibacteria. This analysis revealed that 30.4% (7/23) of the core patescibacterial genera are not represented by the current genome databases **(Table S10).**

**Figure 5.**
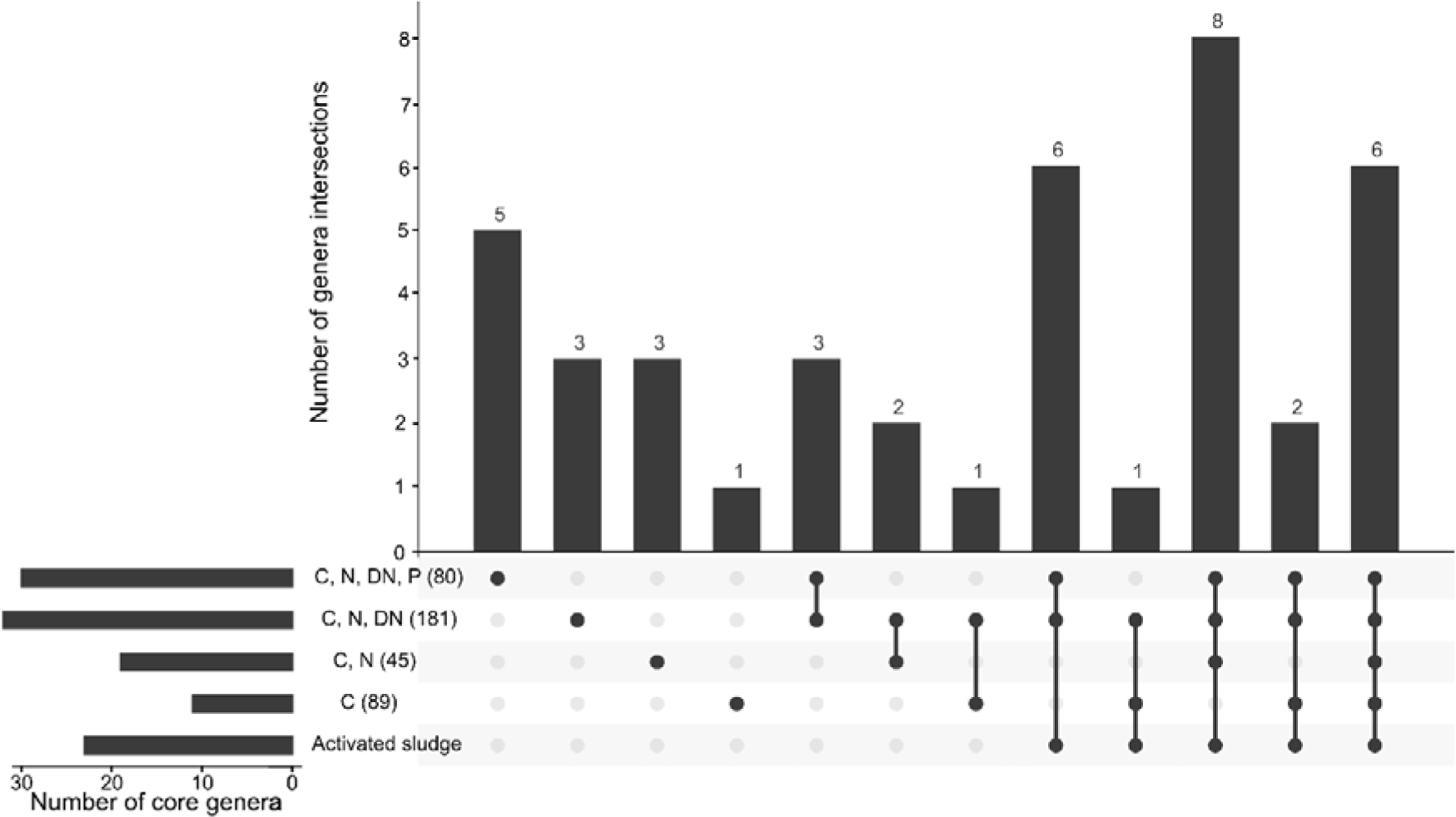
Core patescibacterial genera identified in the four main process types of activated sludge systems. Overlap of process type specific core genera and general activated sludge core genera (identified in the four main process type samples) of Patescibacteria were displayed using a UpSetR plot [31]. The number in brackets after each process type represents the number of samples of each process type used in this analysis.

We further identified patescibacterial core genera for each of the four main process types of the activated sludge systems, using the same thresholds as described above. 11, 19, 32 and 30 genera were identified as core genera for each process type (C / C & N / C & N & DN / C & N & DN & P) respectively **(Table S11-S14)**. In this more refined analysis, one, three, three, and five genera were identified as WWTP-type specific core genera of each process type (C / C, N / C, N, DN / C, N, DN, P) respectively (**Figure 5**). Among these core genera, we found one core genus present in the C, N, DN process type and five core genera specific to EBPR plants exceeding 0.1% abundance in >50% samples, which were identified as “general core” members of these two process types. **(Table S13, Table S14)**. These WWTP function-specific enriched patescibacterial genera may have a potential impact on the respective nutrient removal processes.

We also identified 310 CRAT patescibacterial genera in activated sludge samples. The CRAT genera are not part of the core genera defined above but occasionally show high abundance. While most of the CRAT genera (170) were only detected in one sample with >1% relative abundance, eight genera were detected in more than ten samples with >1% relative abundance **(Table S15)**.

### Potential host-Patescibacteria pairs revealed by co-occurrence network analysis

Patescibacteria have been predicted to lead a symbiotic lifestyle based on their small cell and genome sizes, and their limited metabolic capability [19]. Several studies have successfully applied network-based methods for the inference of symbiotic relationships which were subsequently validated by experimental evidence [33,34]. Here, we performed network analysis at both the ASV and genus levels to answer the questions: (i) which bacteria or archaea are correlated with Patescibacteria in activated sludge systems, (ii) whether different genera of Patescibacteria correlate with the same potential host, and (iii) whether a similar correlation pattern is observed for different ASVs within the same genera.

Genus level network analysis was performed with 395 samples of the four main process types of activated sludge system for which we obtained >5K reads by the modified primer pair. This analysis identified 10 genera of Patescibacteria, all belonging to either activated sludge or process-specific patescibacterial core genera, that significantly correlate with genera from Actinobacteriota, Bacteroidota, and Chloroflexi **(Figure 6).** Although many of these potential hosts belong to uncharacterized taxa (with MiDAS placeholder names), we also identified some potential host taxa for Patescibacteria with important roles in WWTPs. For example, we observed a strong correlation between the genus *Tetrasphaera* and four other genera, including three patescibacterial genera of Saccharimonadia (*Ca.* Saccharimonas, midas_g_67, and midas_g_363) and the genus *Ca.* Accumulibacter. Interestingly, genera midas_g_67 and midas_g_363 are both identified as EBPR specific general core genera (exceeding 0.1% abundance in at least 50% EBPR plants) **(Table S14).** *Tetrasphaera* and *Ca*. Accumulibacter are both abundant polyphosphate accumulating organisms (PAO) in WWTPs across the world, mainly thrive in EBPR plants and are thus expected to be correlated with each other. Another patescibacterial genus - midas_g_2020 of Saccharimonadia - shows an association with the genus midas_g_399 from the class Actinobacteria, which was also found enriched in EBPR plants, and might represent PAO or glycogen-accumulating organisms [35].

**Figure 6.**
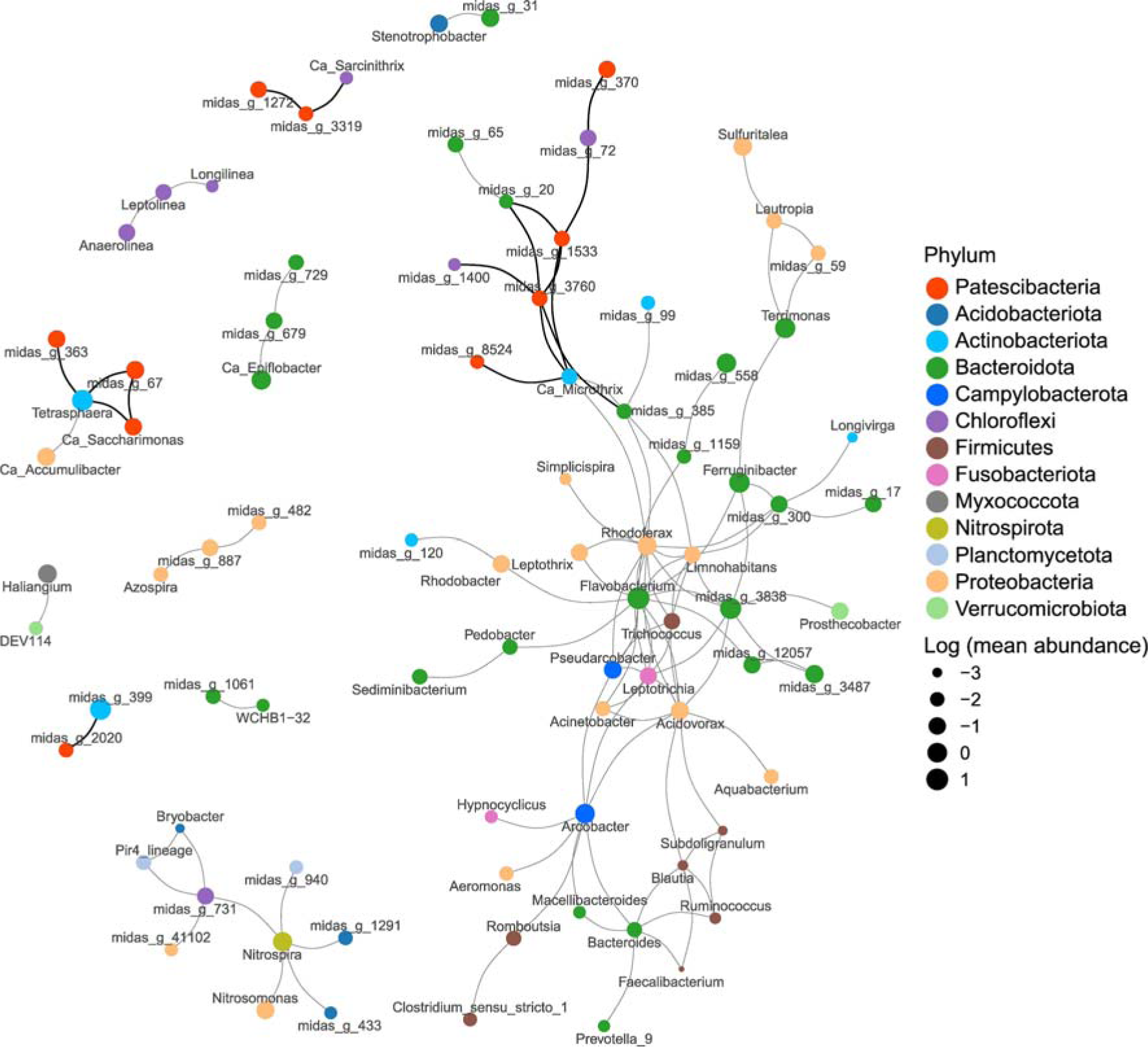
Genus level co-occurrence network reveals potential host-association of Patescibacteria. Genus level network constructed by Sparcc (Friedman and Alm 2012). All genus pairs with a *p* value <0.01 and an absolute correlation value >0.5 are shown in this network. A dark edge color indicates that the edge is connected to a patescibacterial node (red). Size of nodes indicates the mean relative abundance of each genus across activated sludge samples of the four main process types used for the network analysis. The node color indicates different phyla.

Another important group of microbes that was found to be associated with Patescibacteria is filamentous bacteria of the genera *Ca.* Microthrix and *Ca.* Sarcinithrix, the former one known to cause severe foaming problems in WWTPs. In our network analysis, *Ca.* Microthrix is connected with three Saccharimonadia genera (midas_g_3760, midas_g_8524 and midas_g_1533), while *Ca.* Sarcinithrix is connected with midas_g_3319 of the patescibacterial class Microgenomatia. In addition, several correlations between uncharacterized genera and a patescibacterial genus were also detected. For example, midas_g_370 (class Microgenomatia) correlates with midas_g_72 (class Anaerolineae), which is a core member of anaerobic digester communities and enriched in EBPR plants [5].

We further performed an ASV level network analysis to test if the same correlation patterns could be observed **(Supplementary Figure 8),** and explored potential host-symbiont relationships at lower taxonomic levels. Consistent to the network analysis performed at genus level, a correlation was observed between *Tetrasphaera* and members of the Saccharimonadia at the ASV level (two ASVs from midas_s_5 are correlated with two ASVs of midas_s_67 and midas_g_363, respectively). Notably, the midas_s_5 was recently characterised as the most abundant group of PAOs in Danish and global WWTPs and it was proposed to rename it as “*Candidatus* Phosphoribacter” [29]. Furthermore, associations between midas_g_72 and midas_g_370, as well as *Ca.* Microthrix and midas_g_8524 detected in the genus level network were also found in the ASV level network. Additionally, many connections were newly detected, e.g. a connection between ASVs of the genus *Ca.* Villigracilis, which encompasses filament formers that have been proposed to be structurally important for activated sludge flocs [36] and the patescibacterial midas_g_2215 (class Microgenomatia).

## Discussion

Previous global surveys of microbial community composition in WWTPs had been mainly carried out by 16S rRNA gene amplicon sequencing [5,35] and in consequence, the diversity and abundance of Patescibacteria was largely underestimated due to the low primer coverage of this group. Here, we applied a modified V4 targeted primer pair with significantly expanded coverage for Patescibacteria, which revealed a globally high prevalence and relative abundance of this group of putatively host-dependent bacteria in WWTPs. Using the modified V4 targeted primers, we detected an order of magnitude higher relative abundance of Patescibacteria in WWTP samples, elevating Patescibacteria to one of the most abundant phyla observed in these systems. Interestingly, application of our new amplicon primer set also revealed that, in contrast to the previous perception [5], several representatives of the Patescibacteria are members of loose core taxa in microbiomes of activated sludge systems, while other members of this phylum are part of the core community in specific activated sludge processes. High quality metagenome assembled genomes are available for a surprisingly low fraction of these patescibacterial activated sludge core community members. Thus, for many of these lineages, little or nothing is known about their metabolic potential, and their host organisms in the sewage treatment systems remain unidentified. These results, coupled with a significant increase in alpha diversity of detected Patescibacteria and a lack of systematic bias against non-Patescibacteria, suggest that we have developed a primer pair that can be recommended for future analyses also beyond microbial communities in WWTPs for other ecosystems.

In addition to a much better coverage of patescibacterial diversity, the phylum Verrucomicrobiota and Chloroflexi are also poorly covered by the original V4 16S rRNA gene-targeted primers analysed herein **(Table S2, Table S4)**. Importantly, the modifications we made to these V4 primers did not only increase coverage of Patescibacteria, but also significantly increased coverage of members of these clades, resulting in a much better recovery of the abundance and diversity of e.g., members of the genus *Neochlamydia* and several genera of Chloroflexi. It should also be noted that it has been already reported that the original V4 primers can lead to off-target amplification of mitochondrial 16S rRNA genes [37]. While our modified V4 primer pair increased the relative abundance of mitochondrial sequences detected (from 1.4% to 13.2%), this does not speak against using it in environmental or medical microbiome studies, as mitochondrial sequences can be easily removed during amplicon data processing.

In our database comparison, we found that the WWTP-specific MiDAS 16S rRNA gene database performed significantly better than the general SSU rRNA gene SILVA database when used as the reference database for taxonomic classification of WWTP-derived amplicon sequences. This is in agreement with previous findings [5], indicating the MiDAS database should be used as the preferred database for the 16S rRNA gene sequence classification of amplicon datasets from WWTP samples, also when using the modified V4 primers for amplicon generation. Nevertheless, sequencing data generated with the modified primers revealed a noteworthy percentage of novel patescibacterial ASVs that could not be assigned to a genus or could not be mapped to either database using a high identity cutoff, which indicated that there is a considerable diversity of Patescibacteria not currently represented in the MiDAS 4.8.1 database. We interpret the unclassified ASVs as well as the long ASVs detected with the modified primers as detection of true biological novelty rather than sequencing artefacts, since we could map some of the long ASVs to existing patescibacterial MAGs’ 16S rRNA genes, and removed ASVs that could not be classified at phylum level prior to our analyses.

The lifestyle of Patescibacteria remains debated [11,12,19,38]. However, while the host range and lifestyle of major groups of Patescibacteria remain enigmatic, no cultivation-based evidence shows Patescibacteria to live asymbiotically. We applied co-occurrence network analyses to investigate the potential symbiont-host relationships between Patescibacteria and other microorganisms. This network analysis indicated that specific Patescibacteria clades strongly correlate with different groups of filamentous bacteria and PAOs across activated sludge samples. In particular, ASV level and genus level networks yielded consistent results revealing a strong correlation between Saccharimonadia (midas_g_67 and midas_g_363) and the genus *Ca.* Phosphoribacter, suggesting a potential association between some Patescibacteria and microbes that play a key role in EBPR WWTP. Furthermore, strongly supported associations between Patescibacteria and filamentous bacteria in activated sludge (e.g. *Ca.* Microthrix and *Ca. S*arcinithrix) might suggest a similar predator-prey relationship as has recently been described between TM7a and foam-forming *Gordonia* [21]. As some filamentous bacteria cause bulking and foaming problems in activated sludge, some Patescibacteria might in the future even be used for the biocontrol of sludge settling problems and foam formation. In summary, our network correlation analysis provided a candidate list of potential Patescibacteria-host pairs as targets for further ecophysiological studies (e.g., via *in situ* visualisation of both potential partners using FISH).

## Conclusion

Modifications to a primer set targeting the V4 region of the 16S rRNA gene largely improved coverage of Patescibacteria in 16S rRNA gene amplicon sequencing, while being able to detect similar diversity and abundance of other taxa compared to the original primer set. The modified primers can be applied to studies of Patescibacteria in WWTP and other environments, and also improve detection of other taxonomic groups underrepresented due to primer bias (e.g., Chloroflexi). By applying both the modified and the original V4 primer sets to an large collection of global WWTP samples, we revealed an unexpectedly large hidden diversity and abundance of Patescibacteria in different types of WWTPs, and showed that the diversity of Patescibacteria is poorly covered in current databases. We also identified patescibacterial core and CRAT genera in wastewater treatment systems and depicted the distribution of Patescibacteria in WWTPs globally. Co-occurrence network analysis identified potential bacterial hosts that might be associated with Patescibacteria in activated sludge systems. Collectively, these findings demonstrated that the enigmatic Patescibacteria are previously largely overlooked, but are actually highly abundant and diverse players in wastewater treatment microbiomes. Future research should focus on providing metagenomic and ecophysiological insights into all those newly discovered patescibacterial clades that potentially play important roles in different types of activated sludge systems, including the identification of their actual host organisms. If our co-occurrence network-based hypotheses that Patescibacteria interact with selected filamentous bacteria and PAOs as their hosts or prey in activated sludge prove correct, these Patescibacteria may be key to understanding the population dynamics of these functionally important microbial groups in WWTPs.

## Methods

### In silico coverage analysis and modification of 16S rRNA gene-targeted primers

The coverage of 16S rRNA gene-targeted primers **(Table S1)** for amplification of the hypervariable regions V1-V3, V4, V3-V4, V4-V5, respectively, was evaluated on the SILVA database release v138.1 [23] and the MiDAS 4.8.1 full-length 16S rRNA gene database [5] using *in silico* primer match analysis, allowing for zero and one mismatch. Primer match analysis was performed by an in-house script (https://github.com/huiifeng/Patescibacteria_WWTP). The binding regions of primers 515F and 806R of nearly full-length 16S rRNA gene sequences from the MiDAS 4.8.1 database were extracted using another in-house script (https://github.com/huiifeng/Patescibacteria_WWTP). Degeneracy bases were introduced to the existing primer sequences to cover the majority of Patescibacteria sequences in the MiDAS and SILVA databases **(Supplementary Figure 9).** This procedure resulted in modified primer sequences termed 515F_Mod (5’-GTGYCAGMAGBNKCGGTVA-3’), and 806R_Mod (5’-RGACTAMNVRGGTHTCTAAT-3’).

### Sample collection, 16S rRNA gene amplification and amplicon sequencing

Samples and metadata were collected by the MiDAS global consortium (https://www.midasfieldguide.org/global) from 565 WWTPs (one sample per WWTP). Metadata of WWTPs included continent, country, GPS coordinates, sampling date, the temperature in process tanks, process type, and plant type **(Table S7)**. DNA extraction from the WWTP samples was performed by a plate-based extraction protocol using the FastDNA Spin Kit for Soil (MP Biomedicals), as detailed in the MiDAS Field Guide protocols (https://www.midasfieldguide.org/guide/protocols). PCR amplification with the original 515F/806R [6,7] primer set and amplicon barcoding was performed as described in [39]. PCR amplification with the modified primer pair was performed under the following conditions: initial denaturation at 95°C for 5 min; 30 cycles of 95 °C for 40s, 55 °C for 60s, 72 °C for 120s; and a final elongation step at 72°C for 7 min. All amplicons were sequenced on the Illumina MiSeq Platform (v3 chemistry, 600 cycles) as described in [39]

### Amplicon sequence analysis

Raw data processing was performed at the Joint Microbiome Facility of the Medical University of Vienna and the University of Vienna (project ID JMF-2204-03) as described previously [39]. Amplicon sequence variants (ASVs) were inferred by DADA2 package version 1.20.0 [40] applying the recommended workflow (https://f1000research.com/articles/5-1492). Forward and reverse FASTQ reads were trimmed at 220 nt and 150 nt with allowed expected errors of 2, respectively. ASVs were classified with the MiDAS 4.8.1 database and SILVA r138.1 database taxonomy, using the assignTaxonomy function in DADA2 using a confidence threshold of 0.5. ASVs unclassified at the phylum level in the MiDAS database, and ASVs classified as mitochondria or chloroplasts in at least one database were removed prior to downstream analyses.

With the modified primer pair, ASVs classified as mitochondria or chloroplasts accounted for 13.2% of all amplicons across all samples. ASVs that were removed because no phylum classification was obtained accounted for 1% of all amplicons across all samples. With the original primer pair, ASVs classified as mitochondria or chloroplasts accounted for 1.8% of all amplicons across all samples, ASVs removed as unclassified phylum accounted for 0.3% of all amplicons across all samples.

519 WWTP samples for which >5k read pairs were retained after removal of unclassified, mitochondria, and chloroplast ASVs for both primer pairs were selected for primer performance comparison. Alpha diversity of the microbial community was calculated with the ampvis2 package [41] after rarefying sequencing depth to 5,000 reads per sample. Linear regression analysis was done in R 4.1.2 [42]. Differential abundance analysis of results obtained by the two primer pairs was performed by the DESeq2 package [43] using default settings. Representative ASVs of each genus were selected randomly and aligned by Muscle 5.2 [44]. A phylogenetic tree was then constructed by FastTree [45] using -gtr -nt parameters and visualised using ggtree 3.2.1 [46]

Sequence novelty analysis was performed on 519 samples from all process types of activated sludge and other plant types that were deeply sequenced (>5k read pairs) by both primer pairs. BLAST was performed by blastn 2.13.0 [47] with the following parameters: -evalue 1e-5 -num_alignments 1.

In this study, one objective was to reveal potential host association of Patescibacteria in WWTPs. We applied the SparCC (Sparse Correlations for Compositional data) network [48] analysis to both genus and ASVs levels by the fastspar implementation [49] using data obtained from four main process types of activated sludge samples, with 1,000 bootstraps. ASVs not taxonomically classified at the genus level were removed prior to the analysis. In total, 5,860 genera were included in the analysis. The ASV level network was calculated including the 10,000 ASVs with the highest mean relative abundance across the four main process types of sludge samples using 1,000 bootstraps.

## Author contributions

MW, CWH, JMK, PP designed this study. PHN and MKDD provided samples and metadata. HH, BH performed bioinformatic analysis, with input from CWH, PP, MW. HH, CWH, JMK, PP, KK, and MW wrote the manuscript with input from all co-authors and all authors read and approved the final manuscript.

## Acknowledgements

HH was funded by Austrian Science Fund FWF (DOC 69-B). JMK was supported by the Wittgenstein Award of the Austrian Science Fund FWF (Z-383B) to MW. PHN and MKDD were supported by the Villum Foundation (Dark Matter, grant 13351). We are grateful to Jasmin Schwarz and Gudrun Kohl for laboratory assistance with amplicon preparation and sequencing.

## Availability of data and materials

The 16S rRNA gene amplicon sequencing datasets supporting the conclusions of this article are available in the NCBI repository under BioProject accession number PRJNA1013122.

## List of abbreviations

rRNA: Ribosomal ribonucleic acid
WWTPs: Wastewater treatment plants
MiDAS: Microbial database for activated sludge
ASVs: Amplicon sequence variants
GTDB: Genome Taxonomy Database
MBBR: Moving bed bioreactors
MBR: Membrane bioreactor
C: Carbon removal
C, N: Carbon removal with nitrification
C, N, DN: Carbon removal with nitrification and denitrification
C, N, DN, P: Carbon removal with nitrogen removal and enhanced biological phosphorus removal
MAGs: Metagenome-assembled genomes
CRAT: Conditionally rare or abundant taxa
EBPR: Enhanced biological phosphorus removal
PAOs: Polyphosphate accumulating organisms

