## Supplementary Figure for "Global abundance patterns, diversity, and ecology of Patescibacteria in wastewater treatment plants"

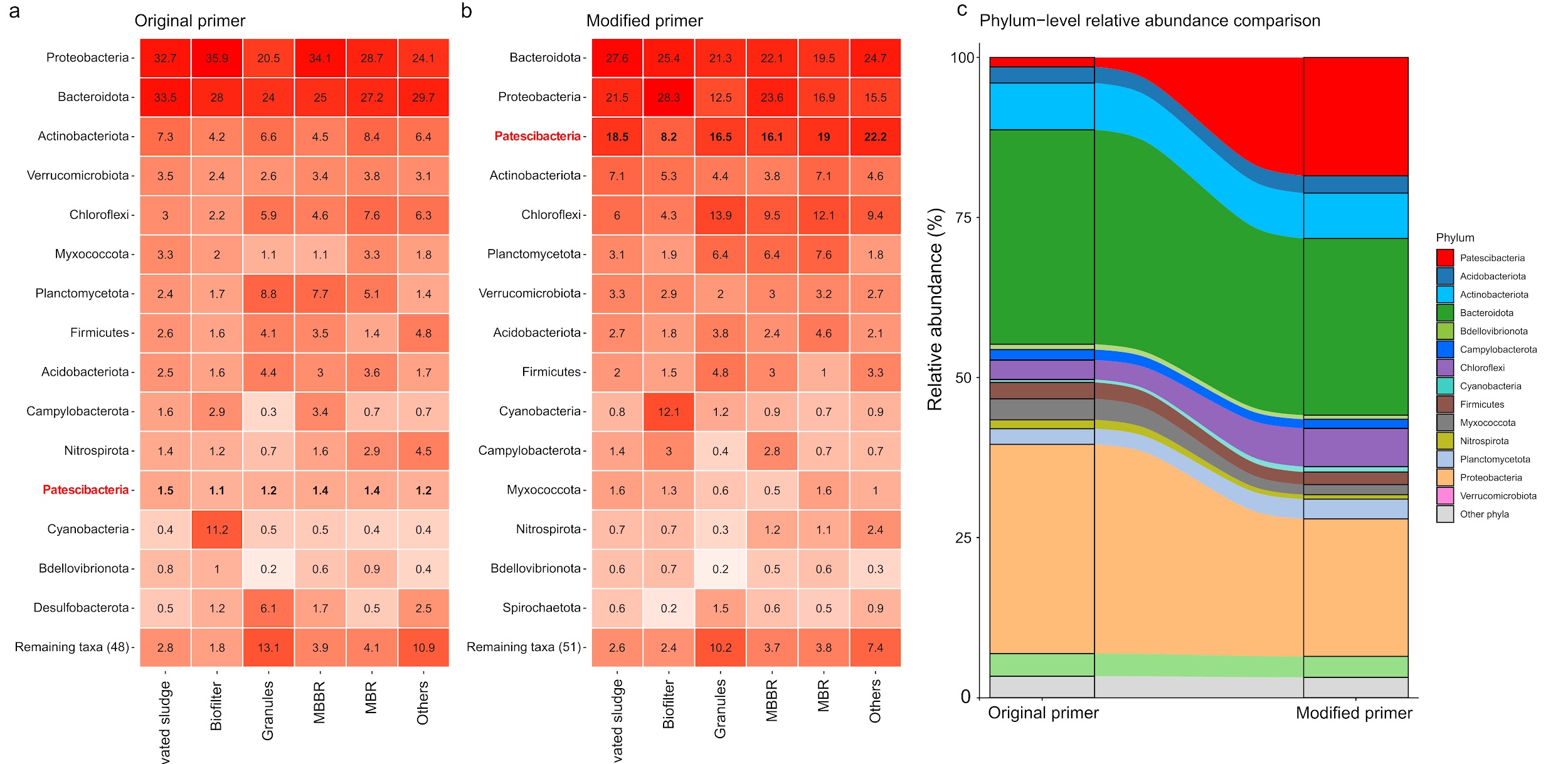

**Supplementary Figure 1. Heatmap of phylum-level abundance grouped by plant type.**

Average abundance of top 15 dominant phyla across different plant types revealed by the original primers (a) and the modified primers (b). The heatmap shows average relative abundance in % for each phylum and plant type. The alluvial-bar plot shows changes in relative abundance of top phyla in the activated sludge systems between original and modified primers (c).

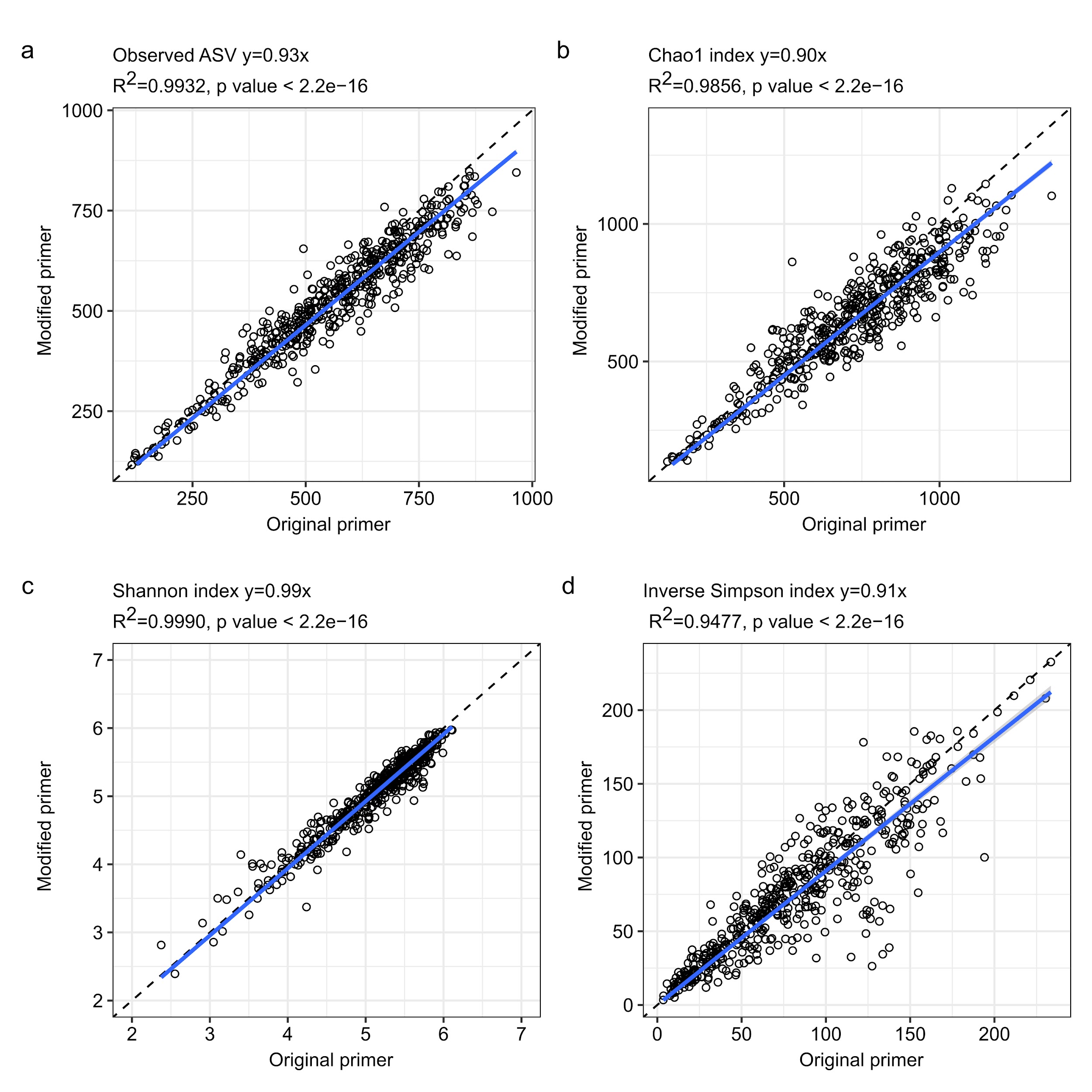

**Supplementary Figure 2. Non-patescibacterial ASV richness comparison between the original and the modified primer pair.**

Regression analysis of alpha diversity indices (ASV richness, chao1 index, Shannon index and inverse Simpson index) obtained by the modified primer pair and the original primer pair of each sample after removing reads classified as Patescibacteria (a-d). Coefficient, R^2^ value and *p* value are indicated in the title of each panel. The observed regression line is depicted in blue, the theoretical 1:1 line is indicated as a dashed line.

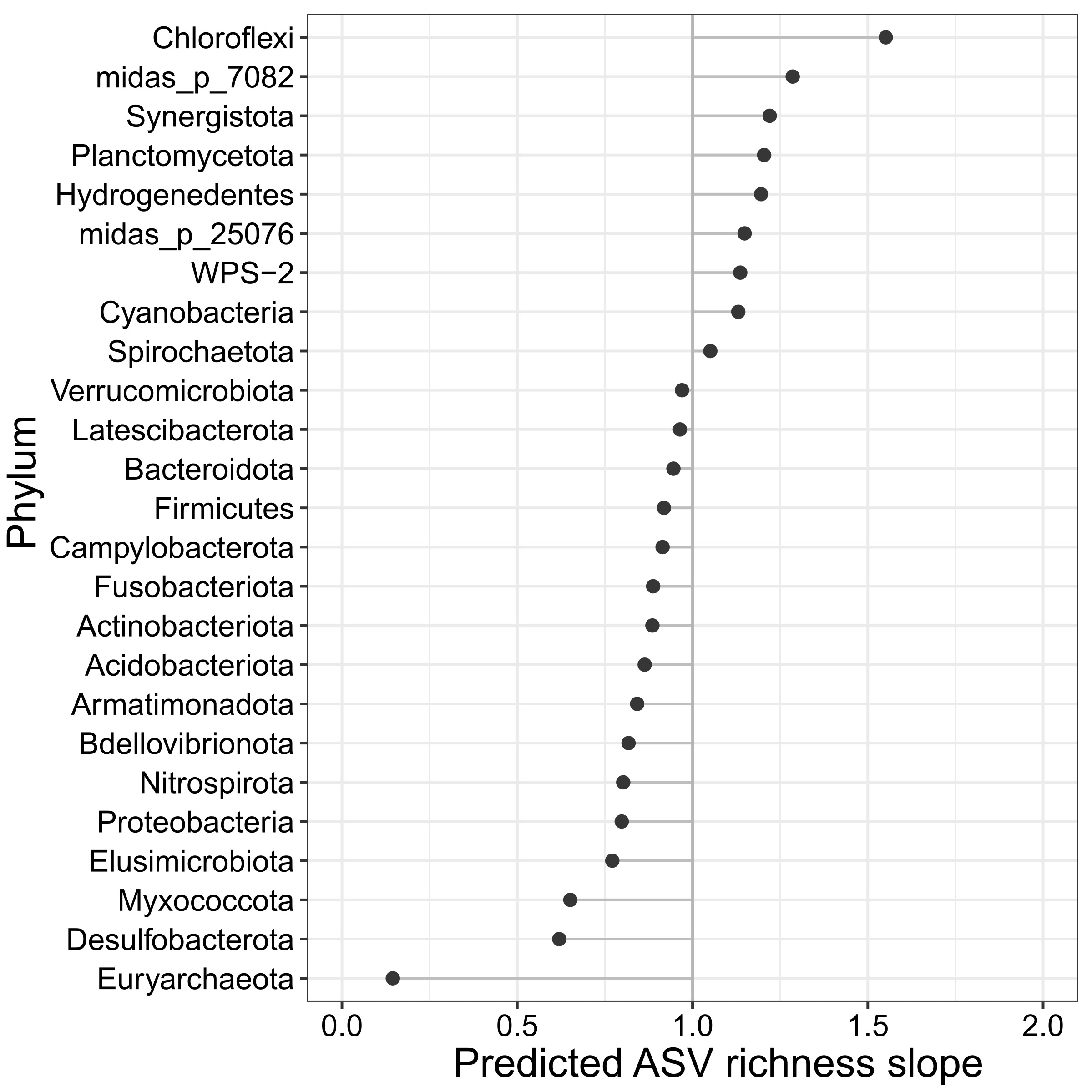

**Supplementary Figure 3. Phylum level richness comparison between the original and the modified primer pair.** The lollipop plot shows for different phyla the predicted ASV richness slope by a linear regression analysis between the ASV richness revealed by the original primer pair and the modified primer pair. A slope of 1 means equal representation of a phylum by both primer pairs. Only phyla with >0.1% average relative abundance in activated sludge samples are shown.

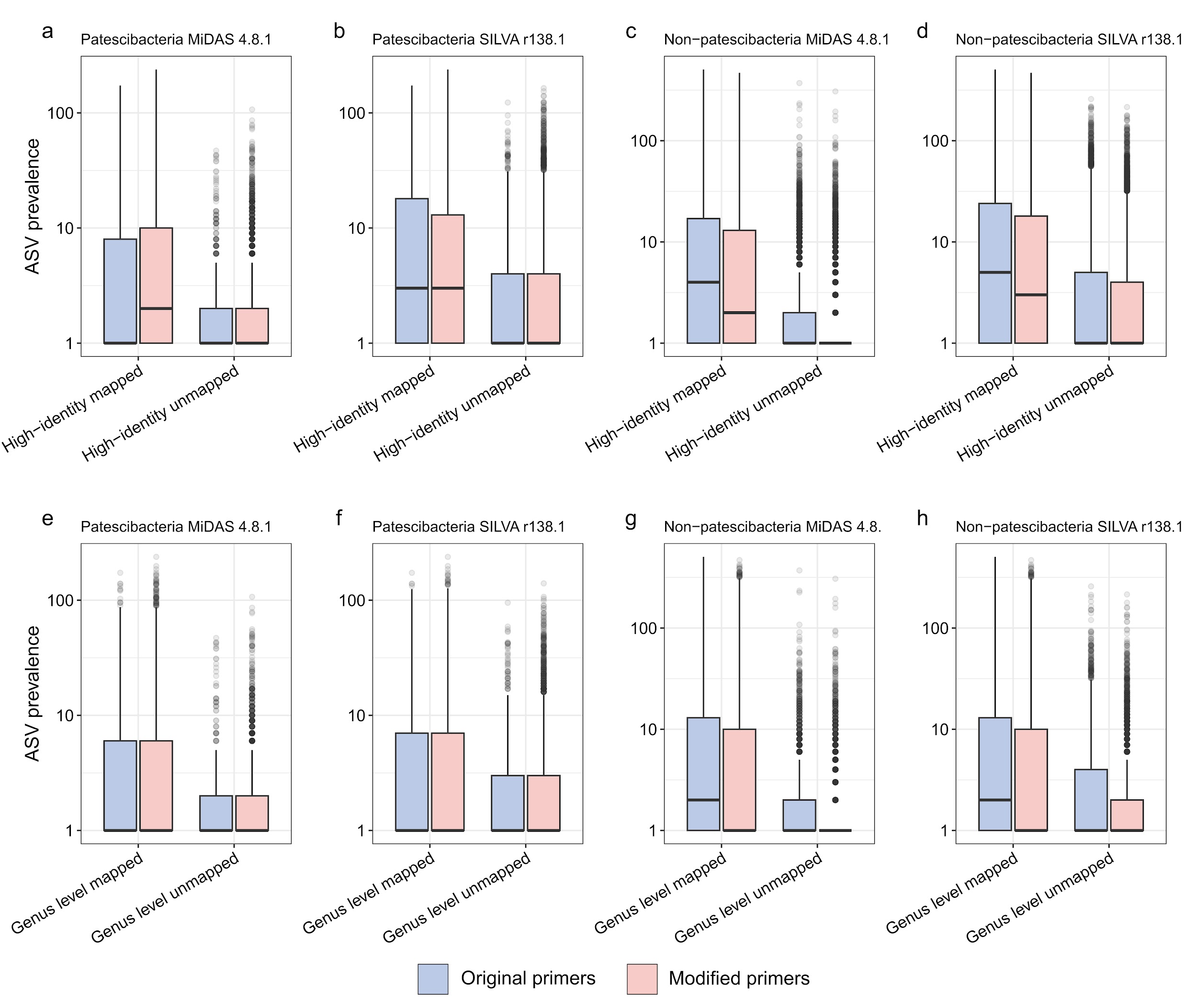
**Supplementary Figure 4. Prevalence of ASVs with and without high-identity (99%) or genus-level identity to reference sequences in the MiDAS and SILVA databases.**

Prevalence of patescibacterial and non-patescibacterial ASVs without high identity (99%) hits (a-d) and without genus level identity (94.5%) hits (e-h) against the MiDAS 4.8.1 and the SILVA r138.1 databases. Y-axis indicates the number of samples where each ASV was detected by the original primer pair and the modified primer pair.

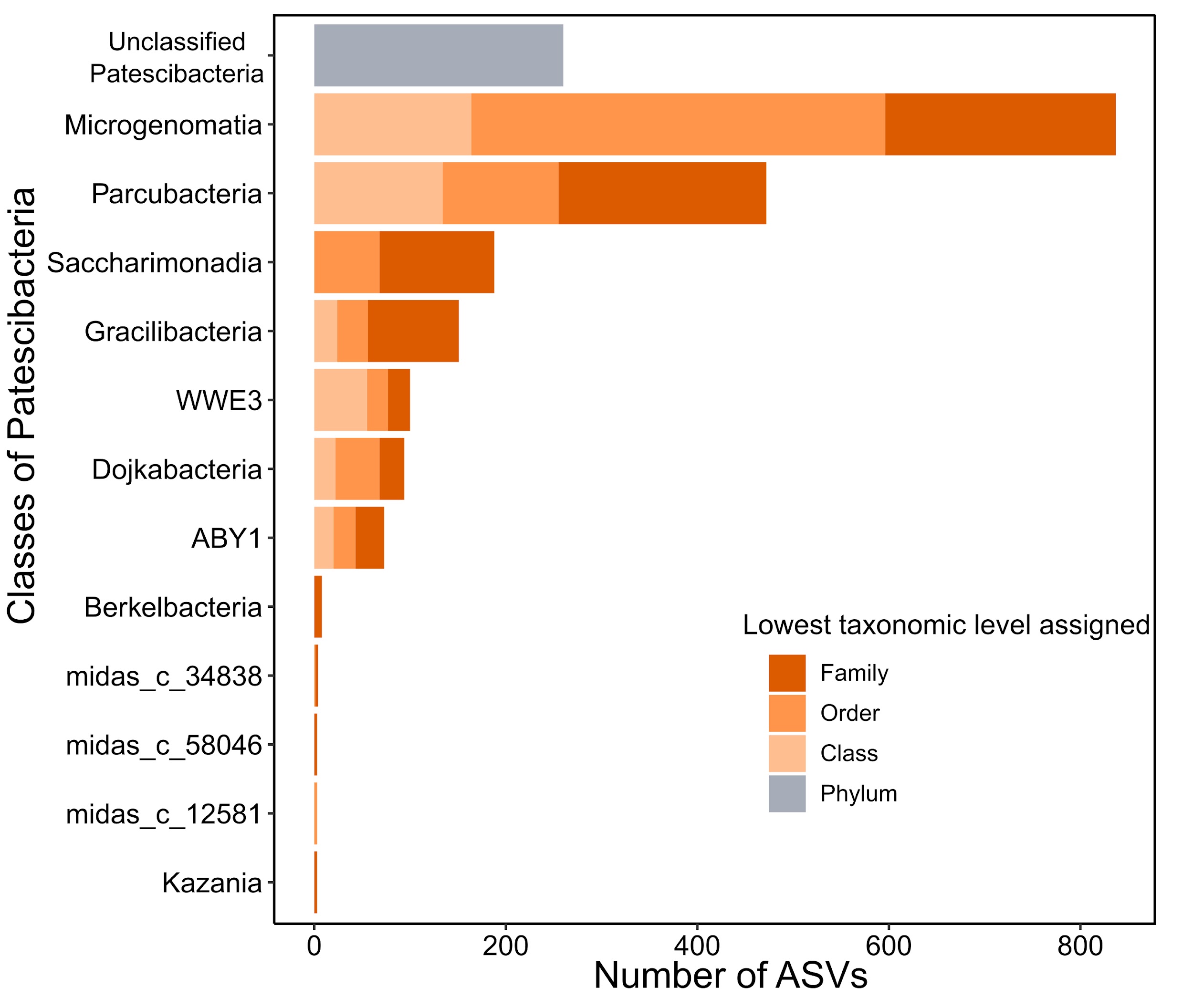

**Supplementary Figure 5. Taxonomic affiliation of unclassified ASVs.**

Number of genus level unclassified ASVs in each class of Patescibacteria (Y-axis). The stacked barplot colours indicate for each Patescibacteria class the number of retrieved ASVs at their annotated taxonomic levels using the assignTaxonomy function in DADA2 against the MiDAS 4.8.1 database.

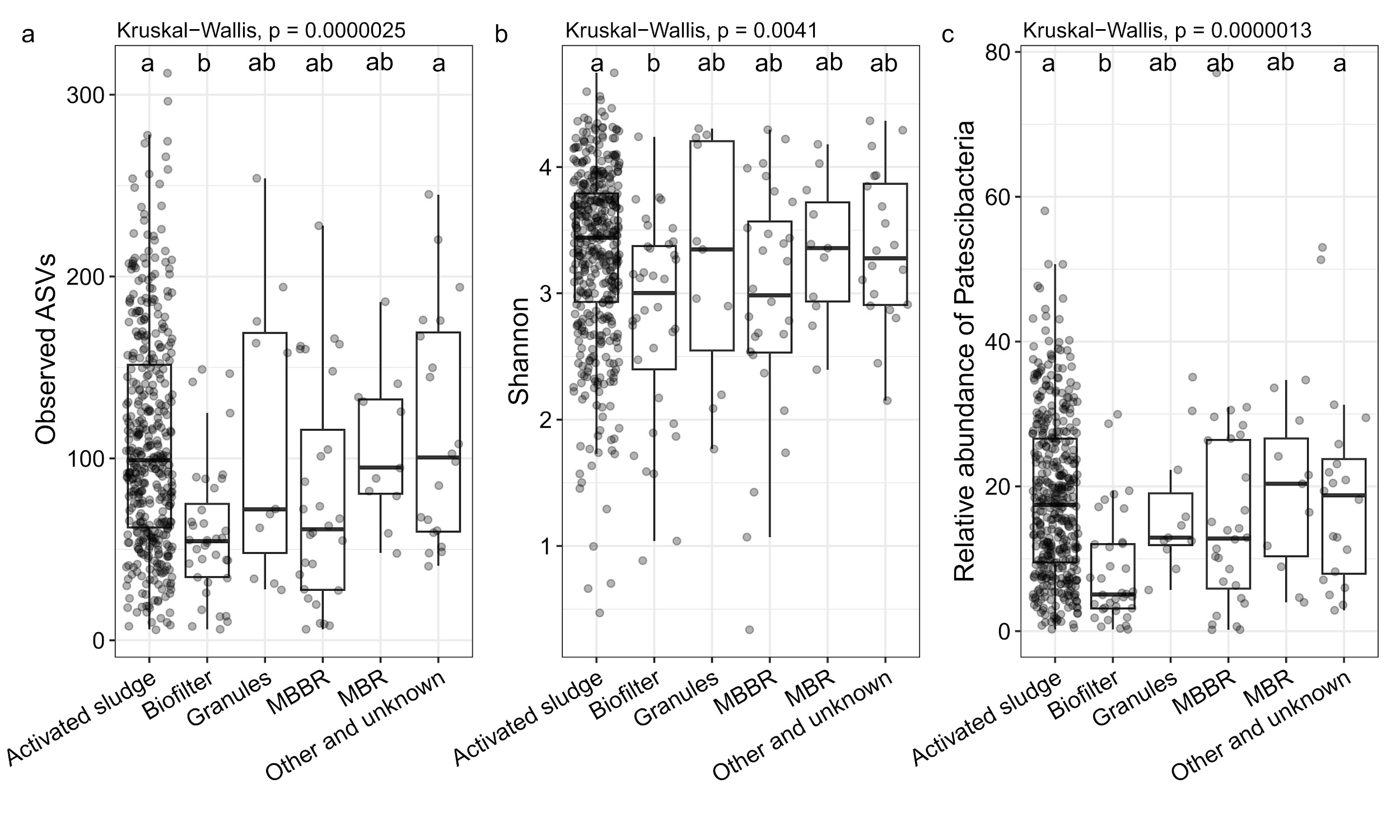

**Supplementary Figure 6. Richness and abundance of Patescibacteria in samples of different WWTP types.**

Observed number of ASVs (a), Shannon index (b) and relative abundance (c) of Patescibacteria in different plant types. The non-parametric Kruskal-Wallis test was used to infer significant differences between plant types. Pairwise comparisons of plant types were done by a post-hoc Dunn’s test (Bonferroni correction, ɑ=0.01) the results are shown with compact letter display (groups that do not share letters are significantly different).

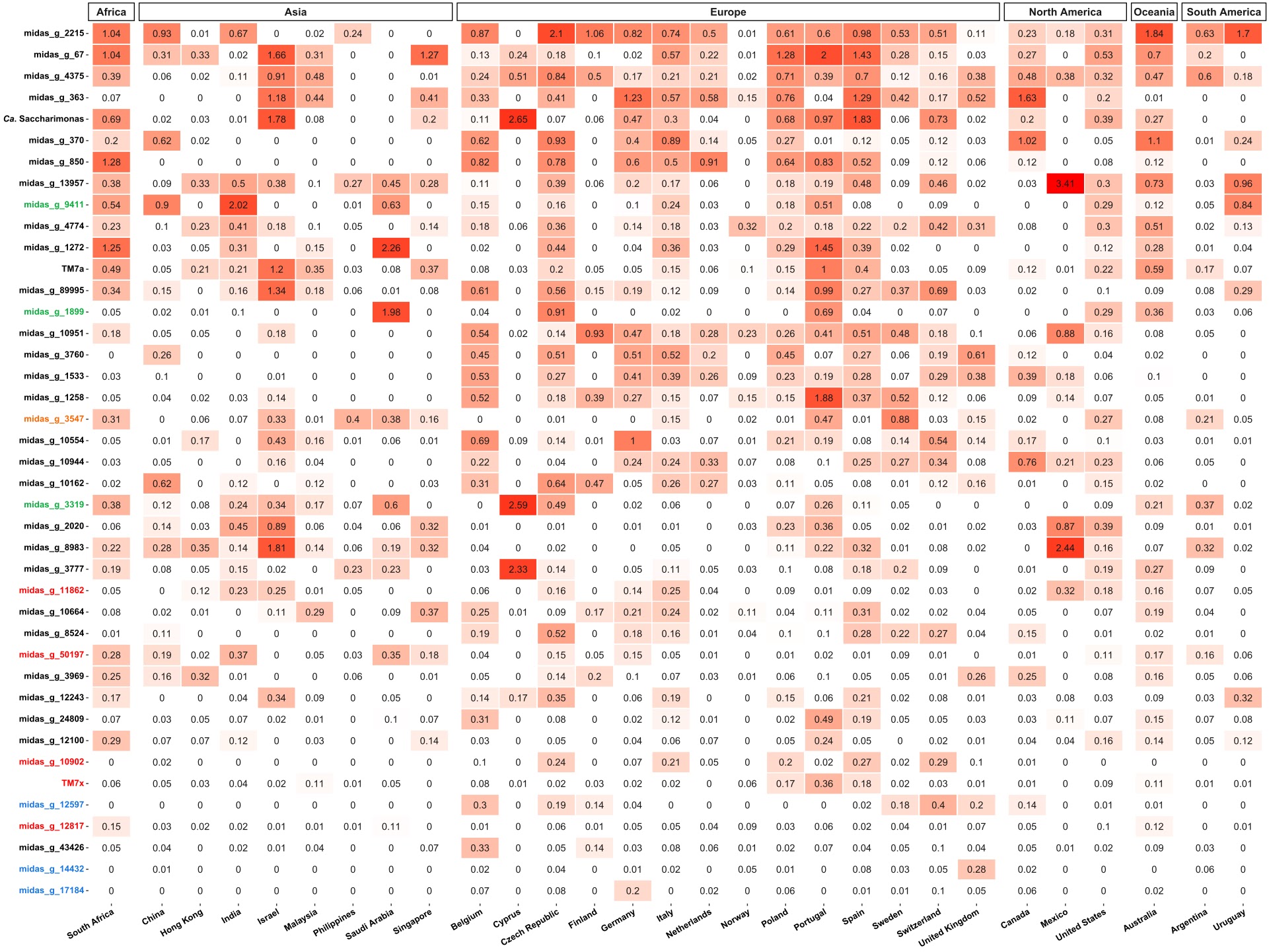

**Supplementary Figure 7. Global distribution of the patescibacterial genera identified as core community members in activated sludge or each process type of activated sludge.**

The heatmap shows average relative abundance of each core genus in WWTPs from each country; countries are grouped by continent. Genus names labelled in orange, blue, green and red represent specific core genera identified in C / C,N / C, N, DN / C, N, DN, P process types, respectively.

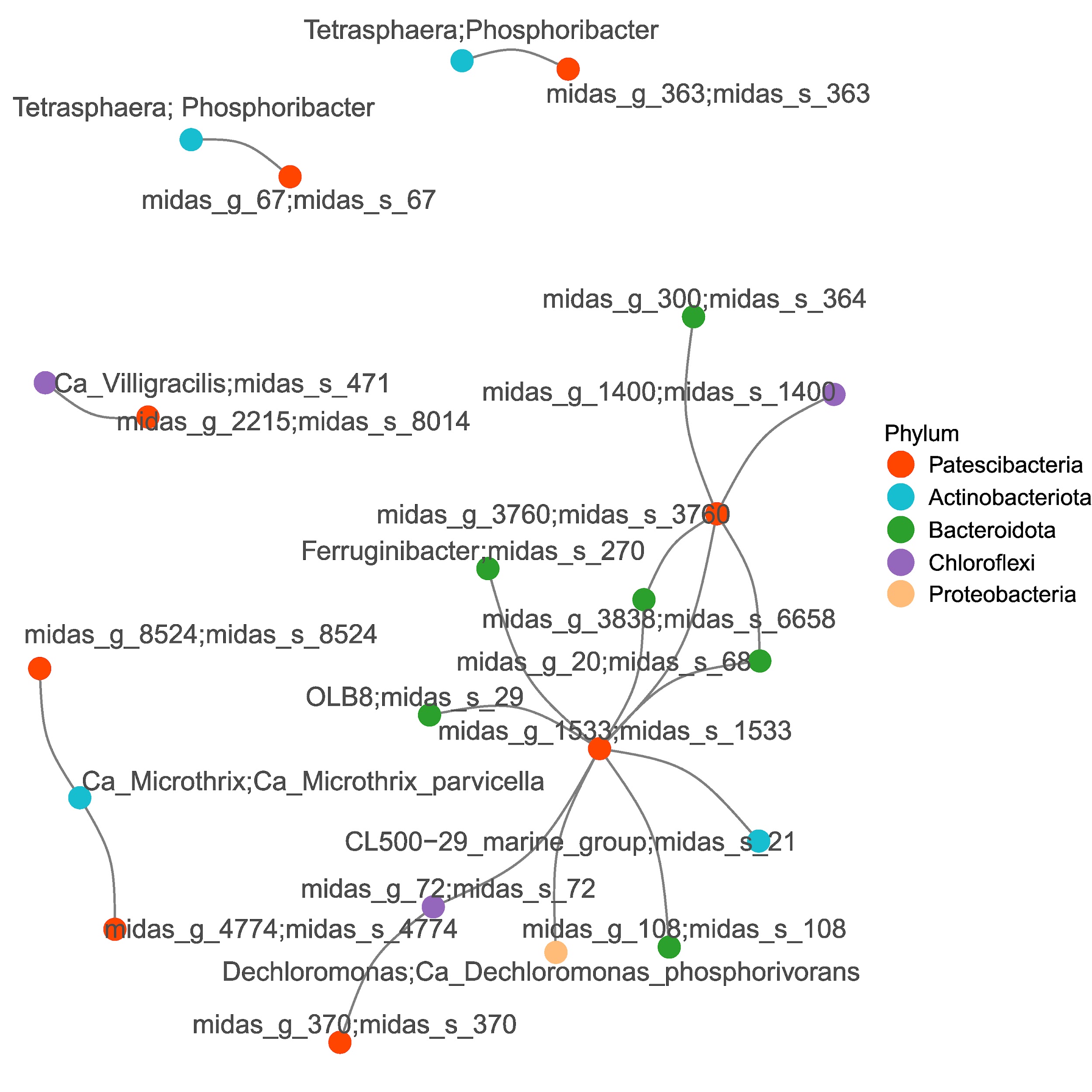

**Supplementary Figure 8. ASV level co-occurrence network revealed potential host-association of Patescibacteria.**

ASV pairs including patescibacterial ASVs with *p* value <0.01 and absolute correlation value >0.5 are shown in this network. Each node represents an ASV, labelled by the species and genus name. The node color indicates different phyla.

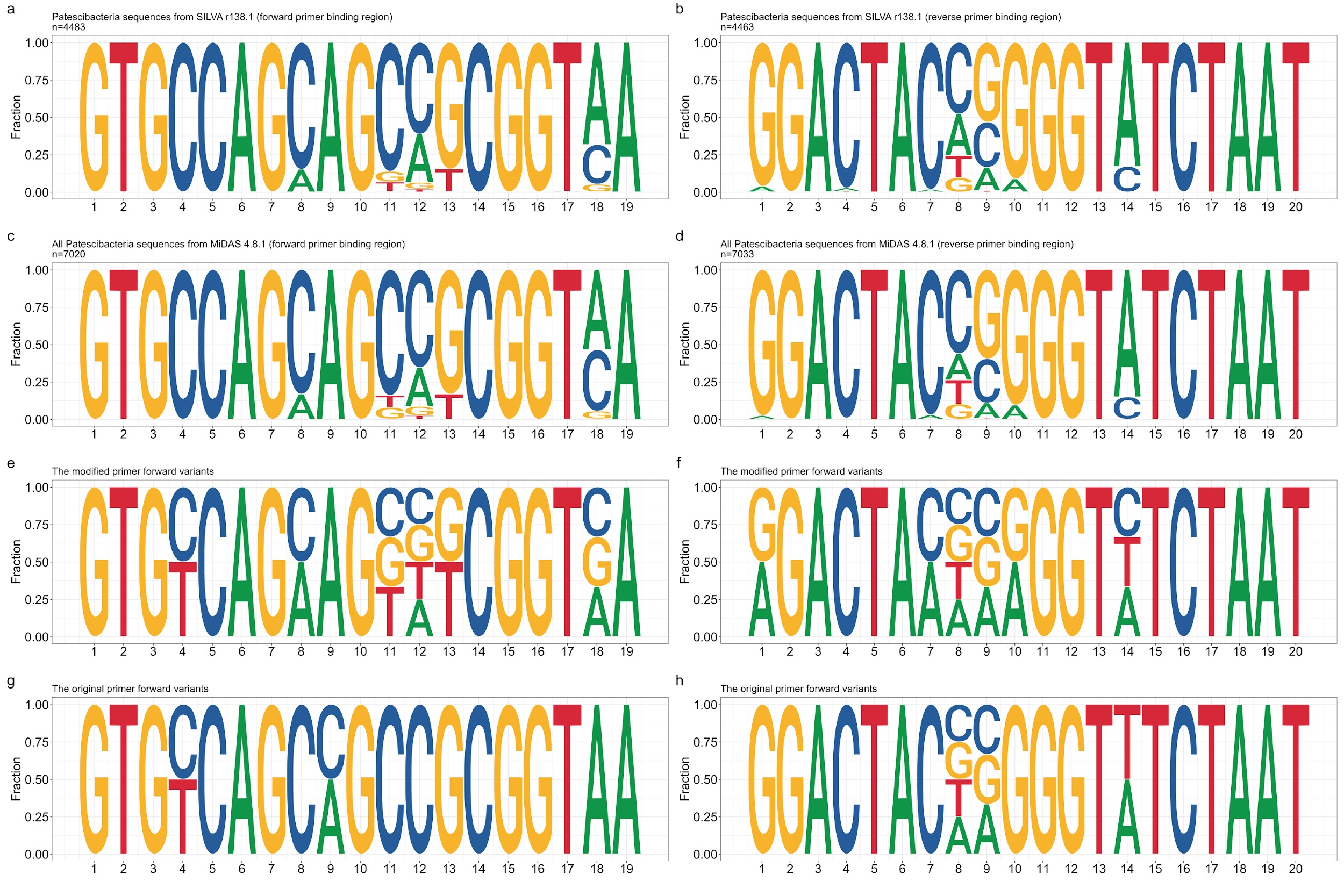

**Supplementary Figure 9. Primer binding region of Patescibacteria 16S rRNA gene.**

Base composition of the binding sites of Patescibacteria 16S rRNA gene sequences for the 515F (positions 515-533)/806R (positions 787-806) primer pair that is widely used for amplicon sequencing. In panel a and b, the results are shown for Patescibacteria 16S rRNA gene sequences retrieved from the MiDAS database (4.8.1). In panel c and d, the results for sequences obtained from the SILVA database (r138.1) are displayed. The modified primer sequence variants are displayed in panel e (forward) and f (reverse). The original primer sequence variants are displayed in panel g (forward) and h (reverse). The Y-axis represents the fraction of A,C,G and T, X-axis represents the position of bases in the primer sequences.

### **Supplementary tables legends**

**Table S1.** Primer sequences used for the *in silico* coverage analyses.

**Table S2.** Coverages of commonly used 16S rRNA gene primers (V1-V3, V3-V4, V4, V4-V5, and modified V4) for different phyla evaluated by the SILVA r138.1 database with zero and one mismatch.

**Table S3.** Coverages of commonly used 16S rRNA gene primers (V1-V3, V3-V4, V4, V4-V5, and modified V4) for different classes of Patescibacteria evaluated by the SILVA r138.1 database with zero and one mismatch.

**Table S4.** Coverages of commonly used 16S rRNA gene primers (V1-V3, V3-V4, V4, V4-V5, and modified V4) for different phyla evaluated by the MiDAS 4.8.1 database with zero and one mismatch.

**Table S5.** Coverages of commonly used 16S rRNA gene primers (V1-V3, V3-V4, V4, V4-V5, and modified V4) for different classes of Patescibacteria evaluated by the MiDAS 4.8.1 database with zero and one mismatch.

**Table S6.** Primer sequences for the modified V4 primer pair compared with the original V4 primer pair

**Table S7.** Metadata for wastewater treatment plants.

**Table S8.** Genus level richness comparisons between the data sets obtained by the original and the modified primer pairs. The slope of the observed ASV and Shannon index were predicted by the linear regression analysis using the modified and original primer sets.

**Table S9.** Blast mapping of patescibacterial ASVs to Danish WWTP metagenomic datasets. Three 100% identity mapping ASVs with >330 base pair coverage were highlighted in yellow.

**Table S10.** Core patescibacterial genera identified if all four main process types of activated sludge samples were analysed together and MAG representation of core genera.

**Table S11.** Core patescibacterial genera identified in activated sludge samples from plants with carbon removal (C).

**Table S12.** Core patescibacterial genera identified in activated sludge samples from plants with carbon removal with nitrification (C, N).

**Table S13.** Core patescibacterial genera identified in activated sludge samples from plants with carbon removal, nitrification and denitrification (C, N, DN).

**Table S14.** Core patescibacterial genera identified in activated sludge samples from plants with carbon removal, nitrogen removal, enhanced biological phosphorus removal (C, N, DN, P / EBPR).

**Table S15.** List of patescibacterial genera that were characterized as CRAT genera

**Table S16.** Correlated genus pairs (X and Y) with p value <0.01 and an absolute correlation value >0.5 identified by network analysis.

**Table S17.** Correlated ASV pairs (X and Y) with p value <0.01 and an absolute correlation value >0.5 identified by network analysis.
